# The transcription factor basal regulatory network of Homo sapiens and Saccharomyces cerevisiae: uncovering the relationship between topology and phenotype

**DOI:** 10.1101/669697

**Authors:** JL Hernández-Domínguez, A. Brass, EM Navarro-López

## Abstract

Transcription factors play a key role in controlling which proteins are made by a cell. As transcription factors are themselves proteins, they are part of a complex interconnected and self-regulated network. We define the transcription factor basal regulatory network (TFBRN) as the network formed by the interactions between transcription factors (TFs) as proteins acting on target genes which are themselves TFs. The question then becomes as to whether topological features of this network are important in determining phenotypes caused by perturbations in TFs. To explore this, we developed two simple TFBRN models; one based on data from human TFs, and the other on the budding yeast. Even from this basic model we did find some very clear correlations between local topological measures and phenotypes seen in cancer and rare genetic diseases. This strongly suggests that the local network architecture of the TFBRN provides important information around the roles of transcription factors and the impacts to an organisation of their perturbation.

**Author Summary:** The human body is controlled by proteins whose production is coordinated by proteins known as transcription factors. These transcription factors can control multiple proteins, including other transcription factors. Does this network itself play any role in determining the properties of the transcription factors and their roles in cancer and disease? In this paper we find that there is a relationship between the local structures in the network and processes such as cancer and rare genetic diseases. We also found a similar relationship between local network characteristics and budding yeast phenotypes. This work therefore shows that simple properties of the network of interactions between transcription factors and their targets can be useful in determining the effects caused by changes in transcription factors (whether through deletion or allelic variation).

## Introduction

All eukaryotic organisms, from the simple baker’s yeast to the whale, have as their building block the cell. This means that many of the same cellular control mechanisms are shared across all eukaryotes. In particular, the mechanisms used to control the processes used to create mRNA and proteins are very similar, a key component in this process are the transcription factors (TFs), which help control the production of mRNA and by association, proteins. They are central to functions like cellular differentiation [1], and how the cell reacts to certain stimulus, like exposure to copper [2]. Transcription factors are proteins that act ultimately to control protein production. As proteins their expression is also controlled by TFs. We are therefore in a position in which TFs can, to a simple approximation, be thought of as a self-regulated network, where TFs controls the production of other TFs. Given the centrality of TFs to all eukaryotic cell function it is interesting to ask whether any features of this network can be used to understand the way in which this network might respond to perturbation, either through deletion of a TF or allelic variation. It might be expected that the sensitivity to such change could be related to phenotypes such as cancer or genetic disease. This work focuses on how the network structure correlates with cancer and other diseases for *Homo sapiens* and phenotypes for the *Saccharomyces cerevisiae*.

Several papers have focused on the behaviour of subnetworks of transcription factors. For example, the network created by TFs Oct4, Sox2, Nang3, STAT3, which have a role in determining pluripotency of the cell, have been studied by Filipczyk et al. [3] in which it showed that the expression of pluripotent factors not necessarily shows differences when under different Nanog levels. A greater understanding on how the TFs pluripotency network works could lead to advances in regenerative medicine [4]. However, such papers have focused only on a subset of the TFs. Less work has been done on understanding the behaviours that could arise from looking at a broader network – one that attempts to capture a more complete set of TF-TF interactions.

We have therefore introduced the idea of the TFs basal regulatory network (TFBRN) – a simple graphical model of the interactions between TFs in the eukaryotic cell. This model is a very simplified model of TFs interactions as it is only concerned to capture cases where a TF is implicated in controlling the expression of another TF. The model does not capture interactions, such as dimerisation, which occurs between TF proteins. The TFBRN is, therefore, a network based only on the interconnection between TFs and the TFs binding sites (TFBS) of genes that are themselves TFs (see figure 1). As such it is a network that captures some features of self-regulation by TFs.

**Figure 1.**
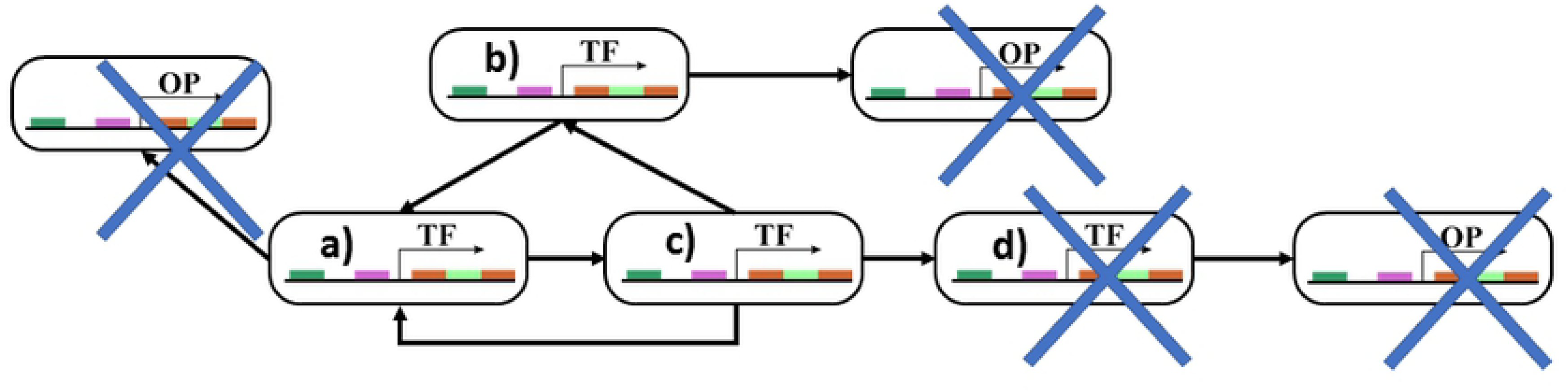
Representation of the general case of the TFBRN. All other proteins (OP) are ignored for this network. TFs a), b) and c) are considered for the network since they control other TFs and are controlled by other TFs. TF d) is not considered for the TFBRN because, although it is controlled by other TF, it does not control other TFs.

Two networks have been constructed based on publicly available information on TFs and TF binding sites, one for the yeast *S. cerevisiae*, and one for a generic human cell. Analyses have then been run in which properties of individual TFs, such as their role in genetic diseases, cancers or cellular phenotype, has been compared against the topological environment within the network. The local topological measures that were included in the analysis include the in-degree, out-degree, eigenvector centrality or eigencentrality of a particular TF (see figure 2). The in-degree and out-degree are measures of the numbers of connections in and out of a node. They can be used to describe and classify a network. The eigencentrality are measures of centrality, or how central (or key) a node is in regard to its neighbours. An interesting feature of the networks constructed from the biological data was the presence of self-loops – TFs involved in controlling their expression. The possible role of such self-loops was also explored. Such analyses have been run on other network types – such as those used to explore relationship between people in social media [5] – but have not been used extensively in the study of TF networks.

**Figure 2.**
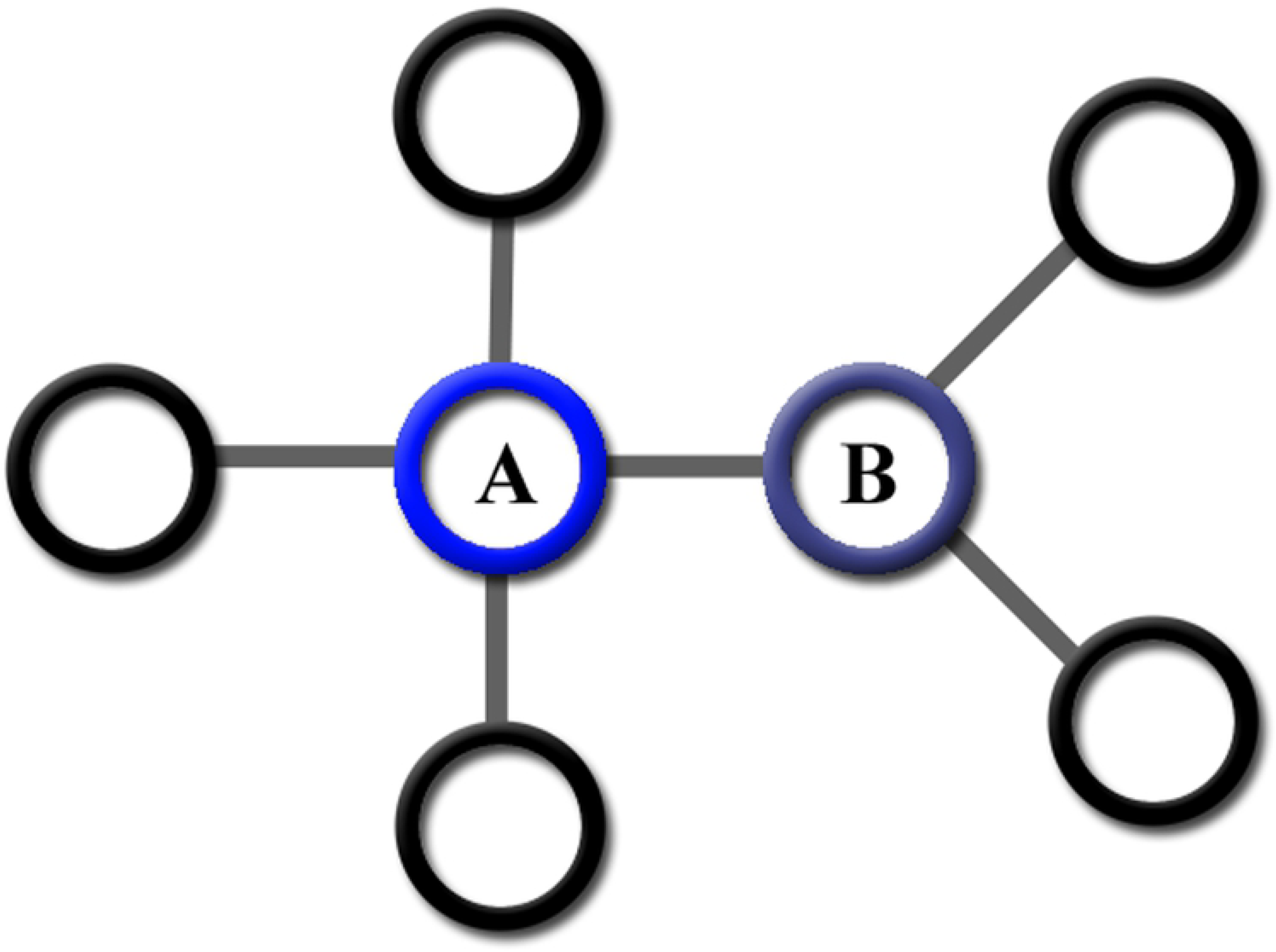
A simplified representation of centrality. Node A) is the most centric node in the network with the highest eigencentrality, followed by node B).

Breitkreutz et al. [6] have looked at the relationship between interaction topology and cancer. However, a key difference with this work is that Breitkreutz et al. focused on molecular signalling network topology and did find it to have a relationship with 5-year survival after diagnosis.

The exploration of the full regulatory network has been done in prokaryotes and lower eukaryotes for the past decades [7,8]. On the other hand, human TF-TF interaction network has been studied generally on particular networks like the Oct4 related networks [9]. Furthermore, works related to the full (or as close as possible) TF-TF interaction network tend to see the agglomeration of networks as a sub-network problem of different cell types [10]. Nevertheless, it is not clear if the works on TF-TF interaction network are done considering just the basal interactions, or they include interaction that ends on TFs that do not interact with other TFs but to “generic” genes.

We are aware that the models contained in this paper capture only a small subset of the transcription control machinery of the cell. The aim of this work is not to precisely simulate a biological system, but to work with a model network based on accurate biological information to ask whether the local TFBRN topology has any predictive power for understanding phenotype. It also ignores the fact that not all TFs are activated at the same time in a cell as DNA histone modification [11,12] can effectively stop the production of specific TFs in specific cell types. However, it is important to try and ensure that the network is based on real data given recent discussions on network topology of biological systems and whether or not they can be modelled as generic scale-free networks [13].

## Materials and Methods

Two transcription factor networks were created, one for *H. sapiens* and one for *S. cerevisiae*. The network is a set of directed links between transcription factors and their cognate transcription factor binding sites on other transcription factors. We chose two databases from which to extract these links, TRANSFAC [14] (version 2013.4) and TRRUST [15,16] (version 2). In TRANSFAC the FACTORS database was queried to that describes the TFs and their target genes. The TRRUST database is a manually-curated database of TFs that have been gathered by sentence-based text mining approach done on PubMed [15,16]. There is some debate as to which proteins can be considered as transcription factors, as discussed in Lambert et al. (27). We therefore looked at another resource, FANTOM5 [17], which is a database of mouse cDNAs and used this as part of a simple bias voting algorithm. This then gave us three resources defining transcription factors. To be classified as a TF in this work the protein would need to be classified as such in at least two of these resources. The *S. cerevisiae’s* TF network was generated from the database YEASTRACT [18], only including as potential interaction those cases for which there was binding or expression evidence.

Under this framework, we could generate a set of links between TFs and their potential binding sites in TF genes. From this set of directed links, we could then generate the TF Basal Regulation Network (TFBRN) adjacency matrix for both the human and the yeast networks. The adjacency matrix is a matrix in which the rows represent the TFs and the columns are the targeted TF genes. By definition, this will be a square matrix as any TF gene is itself a potential target of a TF. Some of these rows and columns could be all 0’s as these were TFs for which we had no data on the TFs that were used to control their expression, or which did not themselves control other TFs. We therefore used a simple algorithm to remove these rows and columns of zeroes from the matrix (Algorithm 1).

### ALGORITHM 1. Cleaning the adjacency matrix to obtain the final TFBRN

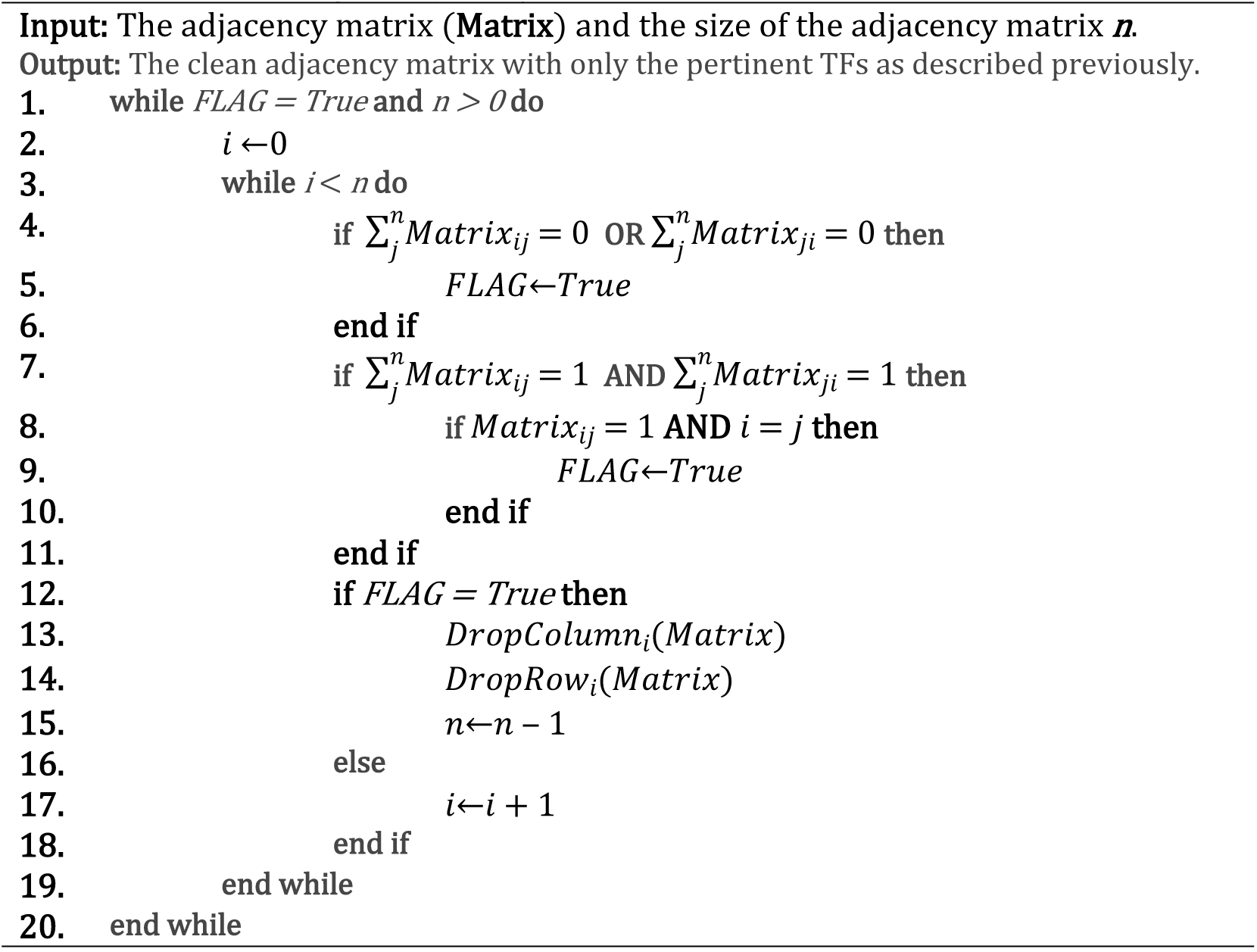

Once the network was cleaned, we proceeded to obtain the network properties. We decided to use the degree of the nodes, the eigen vector centrality or eigencentrality and self-loops. The degree of the node is a measure of how many connections a node has. Since the TFBRN was not a symmetric network, the connections have direction, meaning that there were connections from the node or to the node. The degrees of directed networks are commonly known as in-degree and out-degree. Self-loops, in this work, represent autogenous regulation of gene expression.

Together with the self-loops, the node degrees are some of the few properties that describe the node by itself rather than in relation that the node has with the network [19] or other nodes (e.g. the shortest path length between two nodes). In a sense, it resembles a level of individuality seemingly independent of the network that can be used to explore the diseases or phenotypes of the cell. That is, the minimum basic node properties that can be used to explore the cell phenotypes. The self-loops were represented as a binary variable that describe the presence or absence of self-loops in a TF.

On the other hand, the eigencentrality is a measure of centrality based on the eigenvectors of the network. Centrality, in a broad meaning, is the importance that each node has in relation to the whole network. Because “importance” has different interpretations, there are several types of centralities. From betweenness centrality based on the shortest path length, to the aforementioned eigencentrality, and many others. The objective of the eigencentrality is to score how influential a node is in the network based on the theory that high scoring nodes have a higher impact to nodes directly connected from them than low scoring nodes. That is, a node is more valuable to the network if there are highly valuable nodes directly connected to it. Because of that, eigencentrality is a good contrast to the more individual degree and self-loop properties. The left-eigencentrality is a centrality of a non-symmetric network based on the in-degree, the right-eigencentrality is based on the out-degree.

Cancer is a group of diseases, an abnormal growth, caused by a mutation on the DNA. It is widely known that TFs are an important part of cancer development [20]. We decided to test the TFBRN with cancer given its strong relationship with TFs. To fully explore the relationship between the topological characteristics of the TFBRN and the cancer we divided the cancer list into two lists: oncogenes and antioncogenes. This gave us three groups related to cancer that we named cancer-related (that is if a TF is related to cancer by being an oncogene, antioncogene or both), oncogenes and antioncogenes. The databases used to obtain the oncogenes is ONGene [21], and for the antioncogenes was TSGene [22]. Both databases provide a list that can be easily filtered by the TFs obtained from the process.

On the other hand, diseases in general, though not necessarily related to a mutation on the DNA some of them present a strong phenotype. The phenotypes are linked to genes and, therefore, controlled by TFs. For that, we used a CLIQUE technique that links the phenotypes information available from the Human Phenotype Ontology [23] and the Online Mendelian Inheritance in Man [24]. The results were the number of phenotypes that a gene is linked (the variable named Number of Phenotypes), and the number of diseases related to that gene (Number of Diseases). Furthermore, we created a second binary (yes/no) list that indicates if that gene had related diseases (has-diseases). We reference as VRD (variables related to diseases) to the number of diseases, number of phenotypes and has diseases.

For the *S. cerevisiae* case the decision was to focus solely on phenotypes in general. The yeast case was created as a comparative model for the human network model. We made the decision of using all yeast phenotypes, since most phenotypes could be tracked to one or more genes. We obtained the number of related phenotypes with the database SGD [25], specifically, the YeastMine extracting UI [26].

The random shuffling was done by taking one column, e.g. the in-degree, and permuting the index of the rows. This was done for the degree’s columns and diseases (cancer-related, oncogene, antioncogene, has-diseases) for the human dataset, and this counts as one shuffle. This was done 100,000 times. The same procedure was also done to the yeast dataset. Algorithm 2 shows how it was done for the in-degree and cancer-related including the 100,000 experiments.

### ALGORITHM 2. Random shuffling

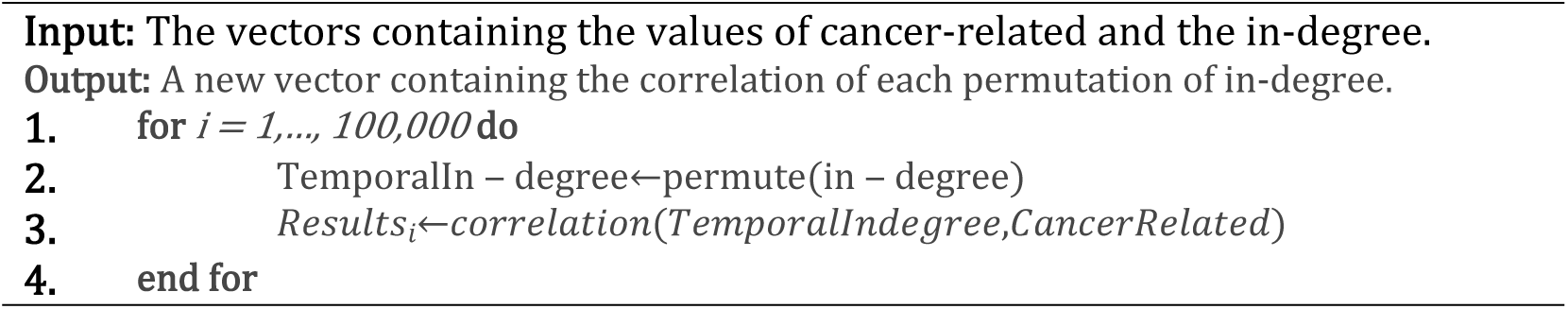

### Software used

The software used to develop the tools to gather the information about the networks and their characteristics are Python (Enthought Canopy Version: 2.1.9.3717 −64 bit-) with the packages NetworkX (1.11-7), Pandas (0.20.3-3), and Numpy (1.13.3-3).

## Results

To find the relationship between network properties and cell phenotypes we need to explore each of the two networks. This would create a base line for comparison between both networks and what the literature says. Once we had explored the networks, we proceeded to seek the relationship between the network properties and cell phenotypes by using Pearson correlation. Even though our main interest in the human relationship with the phenotypes, the yeast provided us with more evidence that this relationship might not be by chance. To further increase the evidence, we developed a randomised shuffle.

The *H. sapiens* TFBRN we obtained from the databases contains 253 nodes. It had an average degree of 4.719 connections (in-degree and out-degree) with a total of 2,386 interactions. It also consisted of 60 nodes with self-loops. Figure 3 represents the *H. sapiens* network. Although both figures (3a and 3c) seems different, they are the same network; Figure 3a highlight the in-degree, while Figure 3c shows the out-degree. The nodes closer to red and larger are the nodes with higher degrees. The degree distribution of the In-degree (Figure 3b) seems to follow that of a scale-free network. The same pattern can be found in the out-degree. Both degree distributions, the one for the in-degree and the one for the out-degree, could be categorised as typical ones of a scale-free. However, number of nodes in the network (only 253) might prove that there was not enough information to categorically establish this fact.

**Figure 3.**
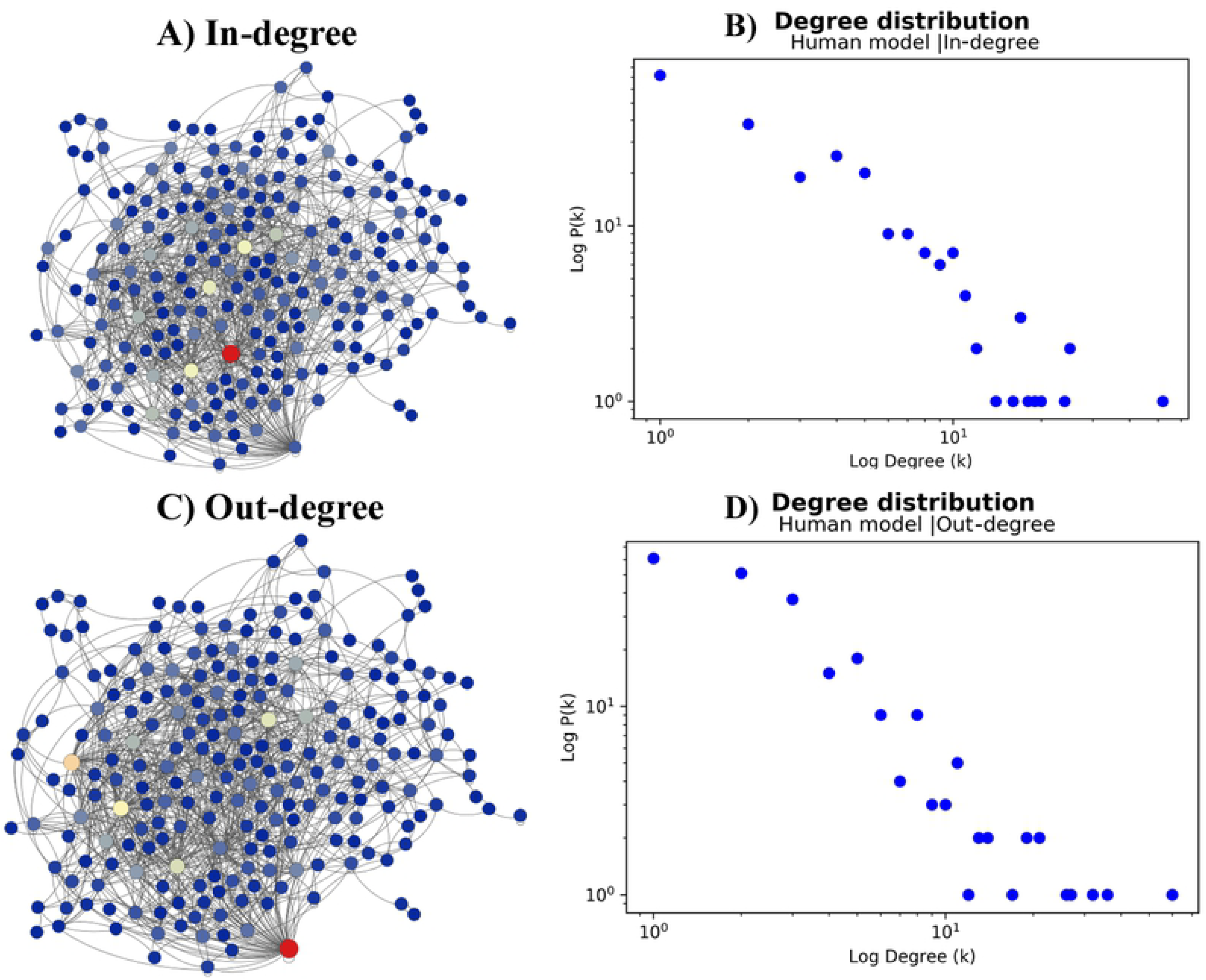
Representation of the TFBRN for the human network. Figure 3a) presents the in-degree value per node. Figure 3b is the degree distribution on a loglog scale and it is close to those of a scale-free network. Figure 3c, on the other hand, is the out-degree.

The *S. cerevisiae* TFBRN, on the other hand, consisted of 154 nodes with an average degree of 26.792 for both the in-degree and the out-degree. It also had 8,252 interactions and a total of 124 self-loops. The number of interactions and self-loops were higher than in the human network. Moreover, the average path length (AVP) is 1.9, which is relatively small for the size of the network and consequently implying the network small-world property. Like Figure 3, Figure 4 size and colour of the nodes represent both degrees and their proportions. Comparing to the human network, the sparsity of the nodes was lower. For the in-degree (Figure 4a) there are fewer dominant nodes overall than for the out-degree (Figure 4c) for the yeast. The Poisson-like distribution of both degrees suggests that the TFBRN for the yeast could be a random-graph network with small-world features or a small-world network.

**Figure 4.**
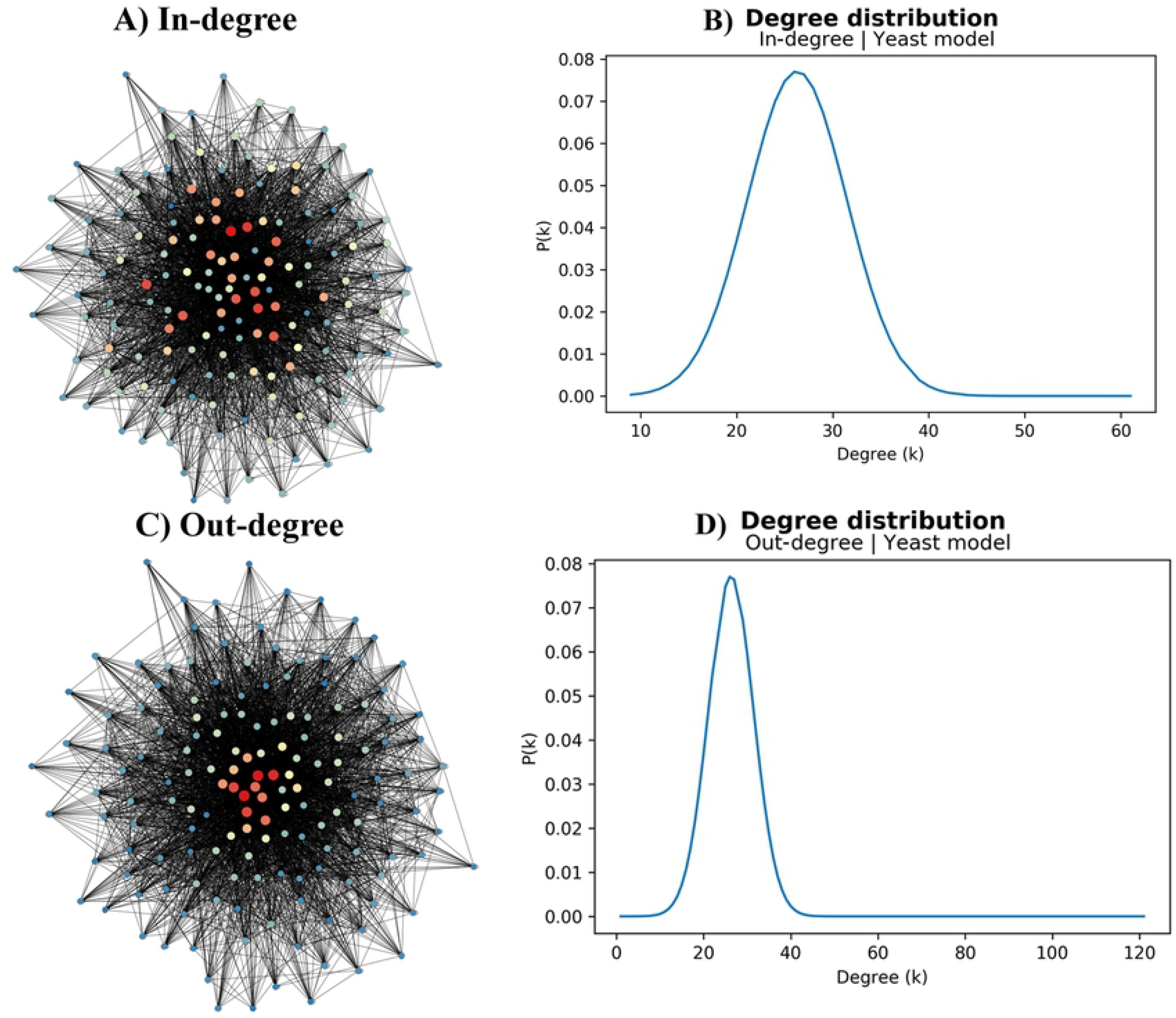
Representation of the TFBRN for the yeast network. Figure 4a represents the in-degree and shows a density that is far superior than the human network. The degree distribution of the in-degree in Figure 4b presents a Poisson-like distribution that is closely related to random and small-world networks. On the other hand, the out-degree presents a higher number of dominant nodes (Figure 4c) and a narrower and more skewed to the left Poisson-like distribution (Figure 4d).

The relationship between the network’s properties and the human cell characteristics is presented in Figure 5. There were four strong relationships with a correlation of an α=0.001 based on Pearson correlation: two in the in-degree and two in the left-eigencentrality. Both network properties had the highest correlation with oncogenes and cancer-related. Only the variables has-diseases and antioncogenes have a medium relationship towards the in-degree and the left-eigencentrality with an α=0.01, respectively. It is important to observe that, although the left-eigencentrality is loosely based on the in-degree, both behave differently with some cell characteristics; antioncogenes and has-diseases are somewhat inverted, while cancer-related and oncogenes are quite similar. Lastly, the least correlated cell characteristics, with an α=0.05, were: antioncogenes with in-degree, has-diseases with left-eigencentrality, cancer related with right-eigencentrality, and oncogenes with self-loops.

**Figure 5.**
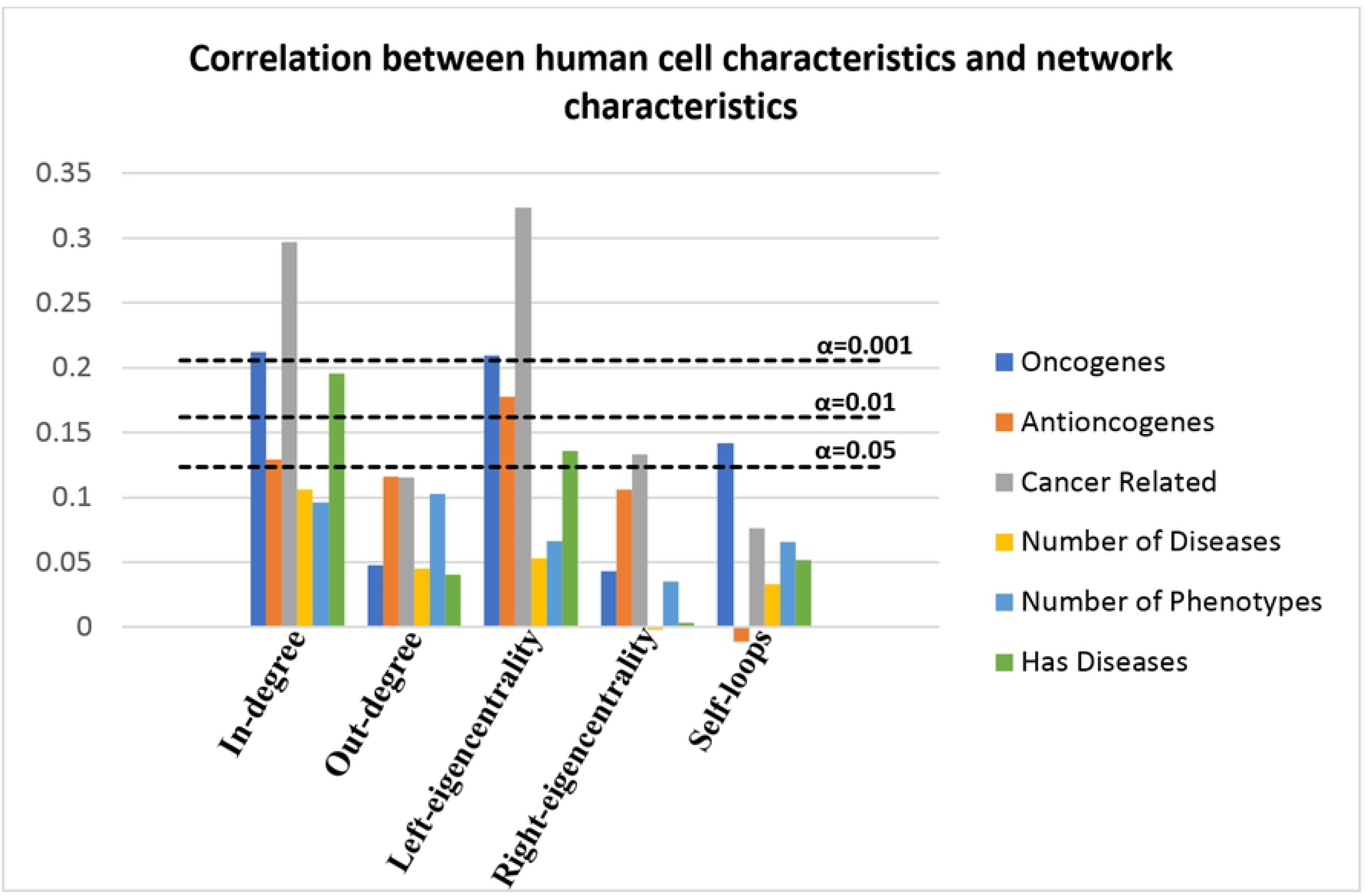
Correlation between VRD and network characteristics. It can be observed a strong relationship between the in-degree and the left-eigencentrality variables and the human cell phenotypes, particularly related to cancer.

While the human’s TFBRN seems to portray the stereotypical scale-free network found on real-world networks, and the topological distribution of the yeast network appears to be close to a random or small-world network, self-loops influence was not widely considered. Works like Lee et al. [27] and MacIsaac et al. [28] indicate that the proportion of self-loops in *S. cerevisiae* were around 10% of the network, much lower than the obtained 80% for the same yeast. This level of auto-regulation is found on prokaryotes like *Escherichia coli* according to Lee et al with a proportion between 52% to 74%. On the other hand, the proportion of self-loops in human’s networks seems to be closer to what Lee et al. and MacIsaac et al. established for yeast.

The relationship between the network’s properties and the human cell characteristics is presented in Figure 5. There were four strong relationships with a correlation of an α=0.001 based on Pearson correlation: two in the in-degree and two in the left-eigencentrality. Both network properties had the highest correlation with oncogenes and cancer-related. Only the variables has-diseases and antioncogenes have a medium relationship towards the in-degree and the left-eigencentrality with an α=0.01, respectively. It is important to observe that, although the left-eigencentrality is loosely based on the in-degree, both behave differently with some cell characteristics; antioncogenes and has-diseases are somewhat inverted, while cancer-related and oncogenes are quite similar. Lastly, the least correlated cell characteristics, with an α=0.05, were: antioncogenes with in-degree, has-diseases with left-eigencentrality, cancer related with right-eigencentrality, and oncogenes with self-loops.

In Figure 6, we present the distribution of the left-eigencentrality and cancer-related, and the ratio of TFs that are considered to have a relationship with cancer. The ratio shows that there are 56% of TFs that are related to cancer by being oncogenes, antioncogenes or both. The pie chart represents the proportion between the TFs that are related to cancer and those that are not related to cancer. That is, any TF that is an oncogene, antioncogene or both is considered a TF that is related to cancer. The boxplot represents the distribution of the left-eigenvalues of both TFs that are and are nor related to cancer. As it can be expected form the correlation, the TFs that are related to cancer have the higher left-eigenvalue value.

**Figure 6.**
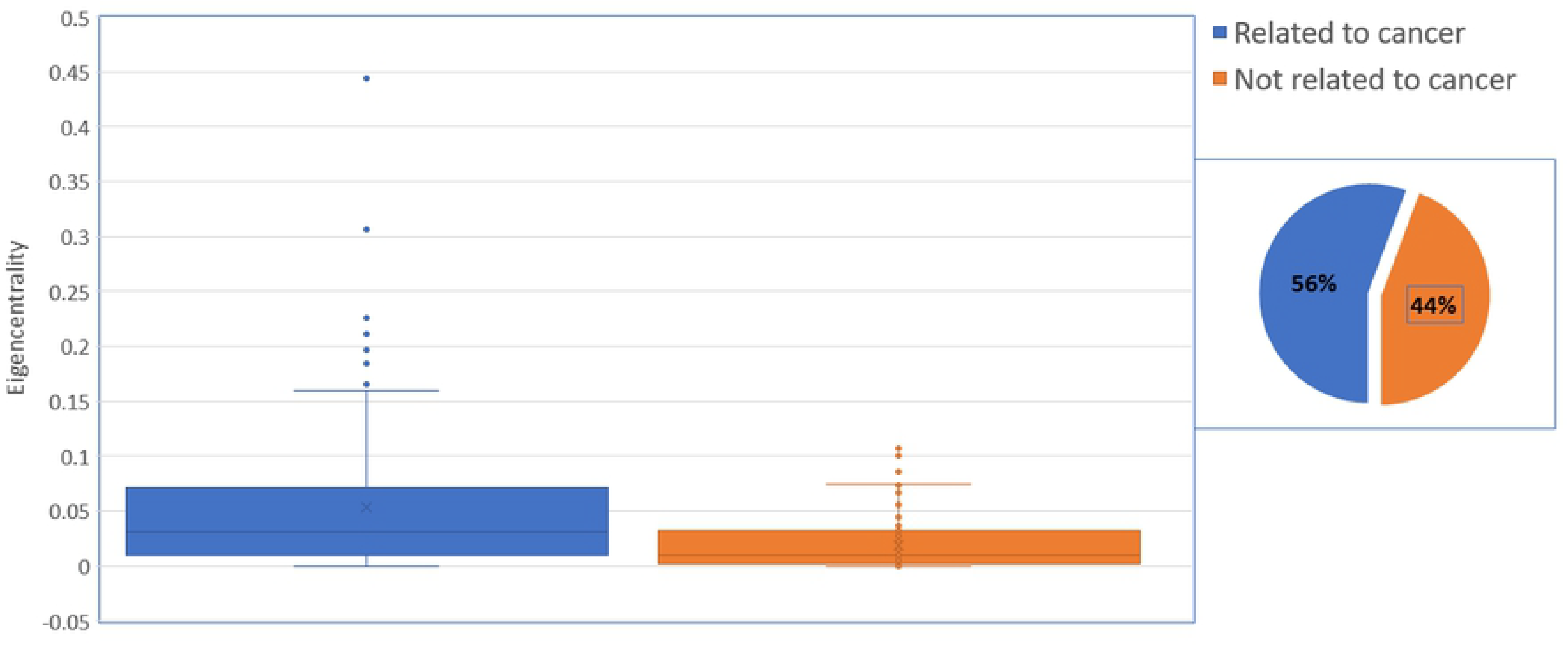
Distribution of the left-eigencentrality and cancer-related, and the TFs values ratio in cancer-relation.

In the yeast, the phenotypes and two network characteristics (right-eigencentrality and out-degree) had not only correlation but a high correlation of an α=0.001 (figure 7). This is in contrast with the highest correlation of the human network properties. The human network properties with the highest correlation were the left-eigencentrality and the in-degree.

**Figure 7.**
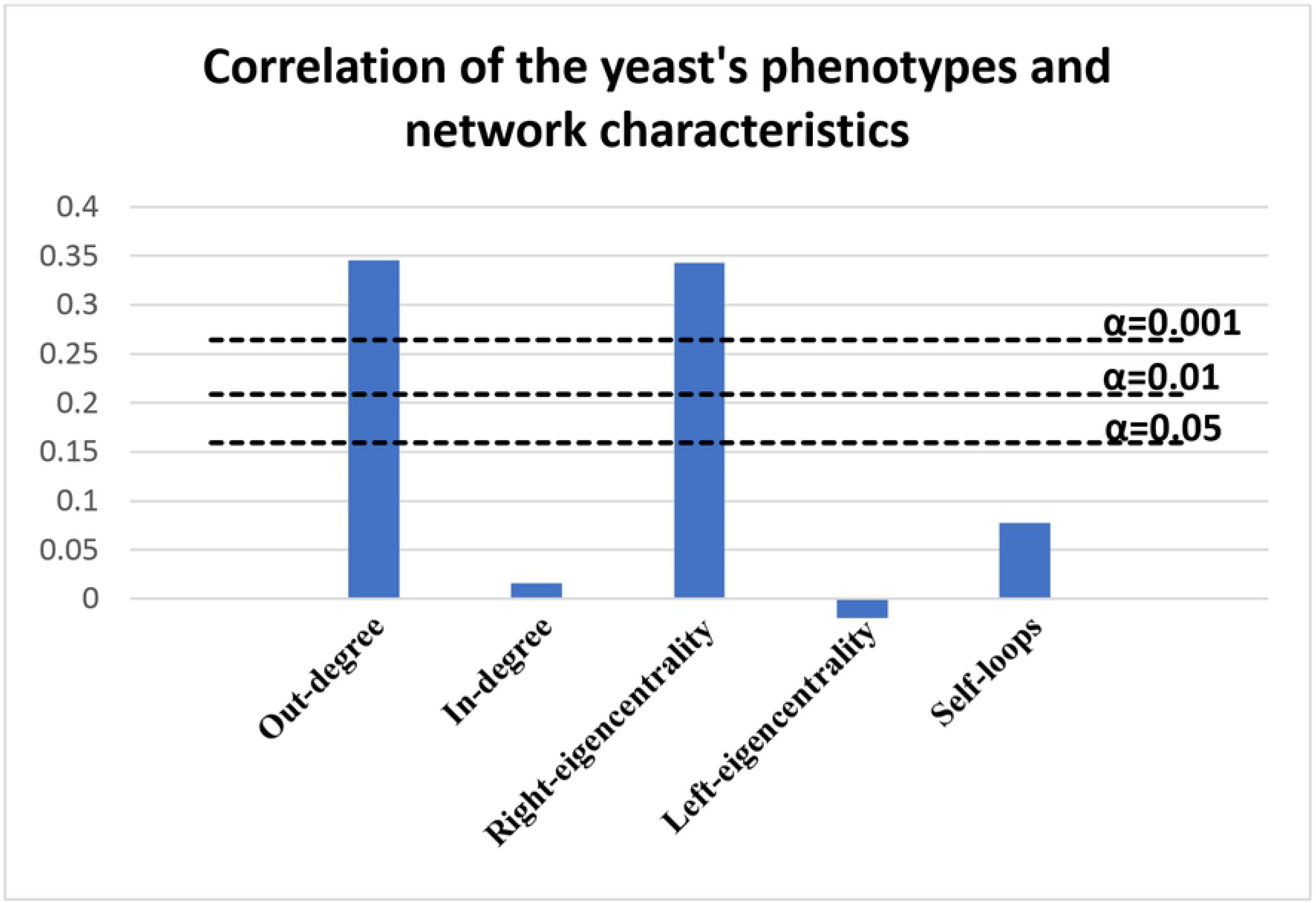
The relationship between the *S. cerevisiae* phenotypes and some network characteristic. The main difference between the human relationships and the yeast is that the yeast strong relationship was related to the out-degree and the right-eigencentrality.

Table 1 presents the results obtained from the randomisation of the values of the network characteristics and diseases for humans related to the in-degree and the out-degree. From the 100,000 random shuffles for the in-degree only the 2.98% had a correlation of 0.05, while 0.65% had a correlation of 0.01 and 0.07% a correlation of 0.001 for the oncogenes. The antioncogenes, cancer-related and has-diseases exhibited results similar to the oncogenes. The out-degree behaves similarly to the in-degree. It is important to note that the previous results the in-degree and out-degree were different (the in-degree had strong correlation with all VRD while out-degree did not).

**Table 1.** The result of the randomised shuffle for the in-degree and the out-degree in the human network.

|  | Correlation | Oncogenes | Antioncogenes | Cancer-Related | Has-diseases |
| --- | --- | --- | --- | --- | --- |
| **IN-DEGREE** | 0.05 | 3.05% | 2.98% | 1.99% | 2.59% |
|  | 0.01 | 0.67% | 0.64% | 0.23% | 0.49% |
|  | 0.001 | 0.05% | 0.07% | 0.01% | 0.03% |
| **OUT-DEGREE** | 0.05 | 2.98% | 2.92% | 2.11% | 2.70% |
|  | 0.01 | 0.65% | 0.63% | 0.28% | 0.49% |
|  | 0.001 | 0.07% | 0.06% | 0.005% | 0.03% |

We replicated the randomisation using the yeast network. The results are shown in Table 2. The results are similar to those in Table 1.

**Table 2.**
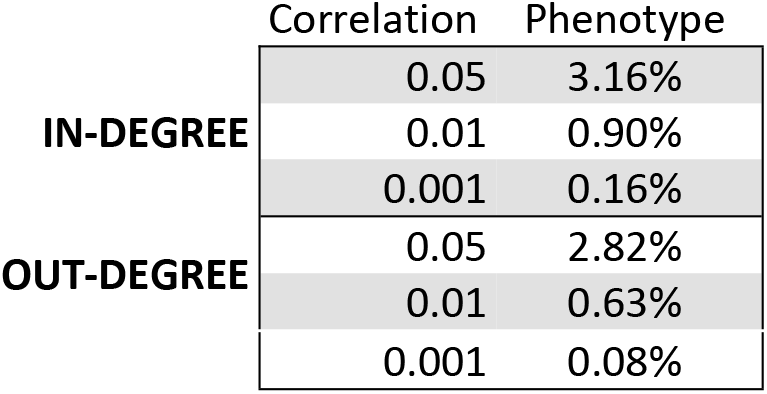
The result of the randomized shuffle for the Out-degree in humans.

## Discussion

We model the transcription factor basal regulatory network (TFBRN) by using multiple databases to explore the possibility that there is a relationship between the topology of the network and associated diseases. The network visualised in Figures 1 and 2 shows a huge topological difference between humans and yeast. The human network has lower average degree than the yeast one, has less self-loops per node and less connections overall, even though it has more nodes. Furthermore, the human TFBRN has characteristics of a scale-free network, while the yeast network is closer to a random or small-world network.

There are a few things to consider when analysing the topological characteristics of both networks. The networks were trimmed to conform the rules stablished for the TFBRN and, therefore, the topological characteristics may not correlate with the usual, non-trimmed, networks. This could potentially explain why the yeast network have characteristics that are closer to a random or small-world network. Another detail is that the network was built with the notion of that all TFs are always active without any difference of cell types (liver cell, brain cell, etc.) or moment in the development cycle of the organisms. These assumptions can lead to a network that deviate widely from the real TFBRN. However, the results may indicate that this may not be the case. Nonetheless, in order to build a network based on biological feature, we used real data with the highest validity that we could find.

We found that there exists a high correlation between the topological characteristics of the TFBRN, cancer and rare genetic diseases for humans, and TFRBN and phenotypes for the yeast. It is also important to highlight that for humans, the dominant correlation was related to the in-degree, while in yeast, it was the out-degree. What is more, the randomise shuffle of the diseases and the network degrees for humans, and phenotype and degrees for yeast, presented a uniform behaviour across all results.

This is significant because, although the dominant degree is different from each species, the behaviour of the correlation was similar. The difference between the human and the yeast degree dominance may be due to that the human has more DNA, thus having more workspace than the yeast. The yeast, by having less space needs to reuse genes looking to reach more genes, or out-degree, while humans seem more focus on having individual genes for particular problems, needing less reach or better gene expression (in-degree).

The correlation found between the topology and diseases and phenotypes, and given the limitations of the study, seems to indicate that even at this level of modelling, there is key information to be obtained. Moreover, the fact that the correlation was highest between cancer than diseases, is interesting. Cancer, by being broadly an alteration of the cell by their genome, seems more related to the topology.

We have considered that it is possible that this correlation was due to randomness rather than actual causality of the mechanics of the cell. And while it is still possible, the results from the random shuffling indicate that this possibility is quite low. Another explanation, at least for the cancer-related correlation, is an over representation in scientific studies around cancer-related genes. This could produce a bias and focus the creation of a topology that correlates with the diseases. This is attenuated by the fact that the *S. cerevisiae* also behaves similarly but with phenotypes.

Aside from the previous mentioned bias, a limitation of the study is that we produce a model based on biology. We overlook several problems in order to make the network. For example, we assume that every single cell type of the human body has the same transcriptional connections. It is highly possible that because methylation each cell has their own transcriptional topology and that some genes are never going to be expressed in an individual cell type. Furthermore, another complication is how complete are the interactions between genes, especially in the human genome. It could be possible that the more interactions are discovered, the correlation might be reduced or even disappear.

## Acknowledgments

The authors want to express gratitude to the Mexican National Council for Science and Technology (CONACyT) for the financial aid and the Wellcome Trust ISSF for the access to the TRANSFAC database.

